# Conserved Non-exonic Elements: A Novel Class of Marker for Phylogenomics

**DOI:** 10.1101/093070

**Authors:** Scott V. Edwards, Alison Cloutier, Allan J. Baker

**Affiliations:** Department of Organismic and Evolutionary Biology and Museum of Comparative Zoology, Harvard University, Cambridge, MA 02138 USA; Department of Natural History, Royal Ontario Museum, Toronto, Ontario, Canada M5S 2C6; Department of Ecology and Evolutionary Biology, University of Toronto, Ontario, Canada M5S3B2

**Keywords:** intron, conserved element, multispecies coalescent, incomplete lineage sorting, biased-gene conversion

## Abstract

Noncoding markers have a particular appeal as tools for phylogenomic analysis because, at least in vertebrates, they appear less subject to strong variation in GC content among lineages. Thus far, ultraconserved elements (UCEs) and introns have been the most widely used noncoding markers. Here we analyze and study the evolutionary properties of a new type of noncoding marker, conserved non-exonic elements (CNEEs), which consists of noncoding elements that are estimated to evolve slower than the neutral rate across a set of species. Although they often include UCEs, CNEEs are distinct from UCEs because they are not ultraconserved, and, most importantly, the core region alone is analyzed, rather than both the core and its flanking regions. Using a data set of 16 birds plus an alligator outgroup, and ~3600 - ~3800 loci per marker type, we found that although CNEEs were less variable than UCEs or introns and in some cases exhibited a slower approach to branch resolution as determined by phylogenomic subsampling, the quality of CNEE alignments was superior to those of the other markers, with fewer gaps and missing species. Phylogenetic resolution using coalescent approaches was comparable among the three marker types, with most nodes being fully and congruently resolved. Comparison of phylogenetic results across the three marker types indicated that one branch, the sister group to the passerine+falcon clade, was resolved differently and with moderate (> 70%) bootstrap support between CNEEs and UCEs or introns. Overall, CNEEs appear to be promising as phylogenomic markers, yielding phylogenetic resolution as high as for UCEs and introns but with fewer gaps, less ambiguity in alignments and with patterns of nucleotide substitution more consistent with the assumptions of commonly used methods of phylogenetic analysis.

---

As a result of advances in DNA sequencing and phylogenetic theory, as well as broader and more aggressive taxon sampling and access to museum specimens, phylogenetics is undergoing a renaissance. “Phylogenomics”, although a term originally coined to denote the increasing need for a phylogenetic perspective when inferring genome function (Eisen 1998), is now meant also to signify the expanded scale in which phylogenetics typically is executed in the era of highthroughput sequencing (Delsuc et al. 2005; Posada 2016). This scaling up has taken two principle forms: increased taxon sampling as a means of producing greater phylogenetic accuracy, and perhaps even more pointedly, increased amounts of sequence data and numbers of loci generated to test a given phylogenetic hypothesis. Many phylogenies now contain hundreds, if not thousands of taxa, although in many cases highly taxon-rich studies still employ a modest number of loci or base pairs in the phylogenetic analysis. Through a variety of next-generation sequencing technologies, systematists now also have access not only to large numbers of loci for phylogenetic analysis but also a wide diversity of genes and noncoding regions for building phylogenetic trees (Bi et al. 2012; Faircloth et al. 2012; Chen et al. 2015). This access to a diversity of loci for building trees has inevitably increased interest in functional ties between phylogeny and genome history, thereby helping re-capture some of the original intent of the term “phylogenomics”. For example, comparison of coding regions generated by transcriptomes across species can reveal key events in the history of adaptation of a clade (Pease et al. 2016), and phylogenetic analyses of conserved noncoding elements and transposable elements in vertebrates has yielded insight into major phases of regulatory evolution (Lowe et al. 2011) and sources of genomic innovation, respectively (Novick et al. 2009).

Despite this progress, in many ways systematists are still constrained by technology in their choice of marker loci for building trees, and this constraint has begun to yield cracks in the vision for phylogenomics going forward (Edwards et al. 2016). For example, transcriptomes are widely used in plant, invertebrate and vertebrate phylogenomics, and with considerable success, in part due to their ease of access in organisms without available genomes and their relative ease of alignment across broad evolutionary distances. Yet, particularly in vertebrate phylogenetics, the deficiencies of coding regions for phylogenetic analysis have long been noted, even in the PCR-era of phylogenetics (Chojnowski et al. 2008; Jarvis et al. 2014). For example, Chojnowski et al. (2008) suggested that introns were superior to coding regions in the phylogenetic analysis of birds in part because of their higher variability. It is also widely recognized that the third positions of codons can become saturated in vertebrate data sets encompassing deeper divergences, sometimes providing unreliable phylogenetic signal. This trend was previously thought to be confined to fast-evolving mitochondrial genes, but is now generally acknowledged for nuclear genes as well, in many cases necessitating removal of 3^rd^ positions of codons or the use of amino acids rather than nucleotides (Cummins and McInerney 2011; Pisani et al. 2015). A compelling example of the challenges of coding regions for phylogenomic analysis has recently been found for birds, where coding regions showed the highest level of among-lineage variation in base composition, resulting in severe challenges for phylogenetic analysis and ultimately yielding gene and species trees with lower congruence than other types of markers (Jarvis et al. 2014). Some of these deficiencies for phylogenetic analysis can be compensated for by improved models of molecular evolution (Philippe et al. 2011; Pisani et al. 2015), partitioning, use of amino acids instead of nucleotides or dropping sites from analysis, yet at the same time there is a clear need for additional kinds of markers that may yield signals more commensurate with the major assumptions of many tree-building algorithms, such as base compositional stationarity.

Ultraconserved elements (UCEs) have also emerged as a major type of marker for phylogenomics, particularly in vertebrates (Faircloth et al. 2012; McCormack et al. 2012; Lemmon and Lemmon 2013; McCormack et al. 2013). These markers, which consist of and whose signal is dominated by the more variable regions flanking highly-conserved core regions, are found throughout vertebrate and other genomes and have a number of features making them attractive for phylogenetics. They are numerous, allowing the accumulation of thousands of markers for a given study, and most importantly, the flanking regions are characterized by high variability, much more so than the conserved regions that are used to identify them. Although this higher variability yields large numbers of informative sites for phylogenetic analysis, it comes at the cost of decreasing reliability of alignments as one moves away from the core, conserved region (Faircloth et al. 2012; McCormack et al. 2013). Perhaps the most useful aspect of UCEs is their convenience: they can be isolated, through hybrid capture or other methods, without knowing anything about the genome of the species under study. In a similar fashion, anchored hybrid enrichment, while not focusing speficially on UCEs, also yields loci easily comparable among genomically novel taxa (Lemmon et al. 2012). Such loci have been readily isolated from hundreds of taxa that are otherwise genomically unstudied. Although many bioinformatics pipelines specifically exclude UCEs that include coding regions, in some studies, UCEs or ‘anchored’ conserved loci include exons (e.g., Lemmon et al. 2012; Prum et al. 2015). Additionally, in several studies not explicitly focused on UCEs, the more variable introns flanking exons have also been accessed in genomically unstudied species in a way similar to the flanking regions of UCEs, using approaches such as exon-capture or anchored enrichment (Lemmon et al. 2012; Hamilton et al. 2016). The convenience of UCEs, transcriptomes and exon-capture when studying organisms whose genomes are not yet sequenced is a major driving force of marker choice in phylogenomics today (Edwards et al. 2016). These markers open up vast areas of biodiversity whose genomes have not yet been sequenced, either due to the unavailability of financial resources, small body size (and hence low DNA yield) of the studied organisms, excessively large or complex genomes, or other factors.

## Conserved non-exonic elements in phylogenomics

Here we analyze a new type of marker for phylogenomics that appears a promising addition to the systematists’ toolkit. Conserved non-exonic elements (CNEEs) are a class of marker that was originally studied in the context of genome function. CNEEs are noncoding regions of the genome that are designated as ‘conserved’ because they evolve slower than a putatively neutral class of sites in the focal clade of organisms (Fig. 1). They are called non-exonic to distinguish them from exons, which also usually evolve more slowly than neutral regions of the genome. CNEEs are distinct from what are often called “conserved noncoding elements” (CNEs) in that they can encompass regions of the genome that are sometimes transcribed but are not in exons. It is often unclear whether genomic regions are transcribed and made into proteins, and recent work from the ENCODE and other studies suggests that many regions previously thought to not encode proteins may in fact be transcribed and translated (Ji et al. 2015). Like UCEs, CNEEs have in many cases been found to act as regulatory enhancers, recruiting transcription factors to influence the expression of nearby or distant genes (Kvon et al. 2016; Leal and Cohn 2016). Crucially, however (Table 1), CNEEs differ from UCEs in not being “ultra-conserved”: whereas the core regions of UCEs are often identified on the basis of >95% or higher sequence identify between genomes, the core regions of CNEEs are designated as conserved only because they evolve more slowly than a putatively neutral rate. As a result, CNEEs often exhibit moderate levels of variability, especially when compared to the core regions of UCEs (Siepel et al. 2005a). This tendency raises the possibility that CNEEs might contain sufficient variability to be useful in phylogenetics, while at the same time exhibiting alignments of a quality that matches or exceeds those of the flanking regions of UCEs or transcriptomes.

**Figure 1.**
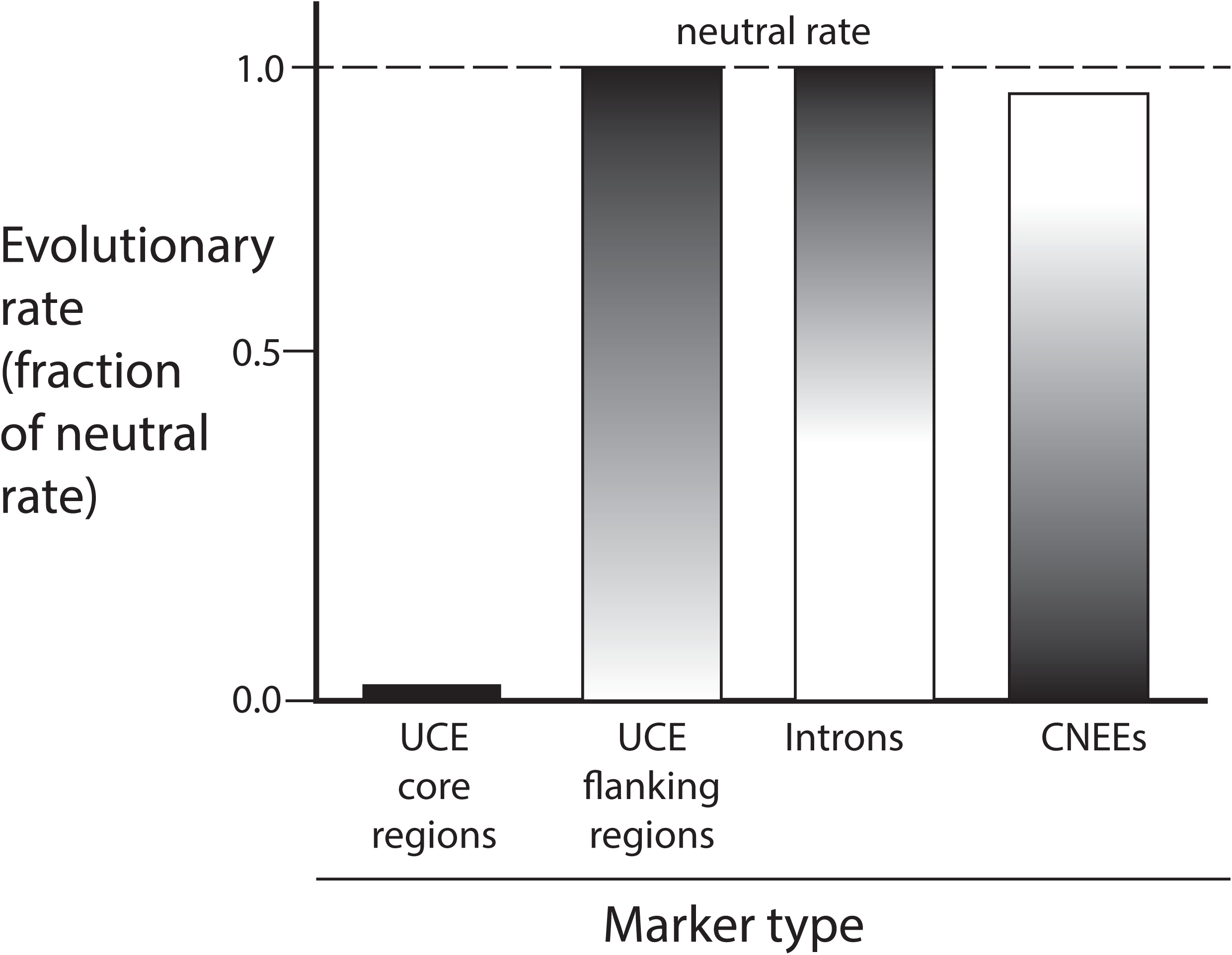
Hypothetical schematic of comparisons of evolutionary rates of different non-coding markers discussed in this paper. The shading of each bar is meant to indicate the distribution of rates within the range indicated by each bar. Thus, as found in this paper, introns are somewhat more variable than UCEs, and CNEEs are less conserved than classically defined core regions of UCEs but not as variable as introns or the neutral rate.

**Table 1.** Differences between vertebrate CNEEs and UCEs for phylogenetic analysis. This table focuses on the classical definition of UCEs as originally described by Bejerano et al. (2004) and Faircloth et al. (2012).

| UCEs | CNEEs |
| --- | --- |
| Individual elements found throughout vertebrates, from fish to mammals/birds | CNEEs arise variably in evolution, and are not necessarily found in all vertebrate clades |
| Information in flanking regions principally used for phylogenetic analysis | Core region used for phylogenetic analysis |
| Can be coding or noncoding | Only noncoding |
| Core regions are ultraconserved | Core regions evolve slower than neutral regions, and thus can evolve faster than UCE core regions |
| Discovered via arbitrary conservation and length thresholds on syntenic-aware whole-genome alignments | Discovered via tuned hidden Markov model |

CNEEs also differ in the means by which they are identified in genomes. UCEs were initially identified using arbitrary thresholds of conservation and minimum length applied to synteny-aware whole genome alignments of a few exemplar taxa; they are often localized in additional genomes by blast searches using previously identified UCEs (McCormack et al. 2012). By contrast, CNEEs are delimited using statistical approaches, such as hidden Markov models (HMMs), wherein the rate of candidate genomic region is compared to the rate at a class of putatively neutral sites (Siepel et al. 2005b, Table 1). These models are usually applied to the entire set of species under analysis (although here we use a hybrid approach in which vertebrate CNEEs previously identified using a set of aligned vertebrate genomes are identified by blast in additional bird genomes). Fourfold degenerate sites of protein-coding genes are the most commonly used class of site to generate a baseline pattern of substitution. (It is reasonable to question whether 4-fold degenerate sites in coding regions are genuinely neutral and alternatives, such as ancient transposable elements that do not appear to have assumed functions, have been used as the putatively neutral class (Siepel et al. 2005a)) Although CNEEs do overlap with the core regions of UCEs in genomes, and include all noncoding UCEs in principle, the use of only the conserved core region of CNEEs distinguishes this class of marker and ensures that the sequences we use here for phylogenetics do not fully overlap with those of UCEs. Additionally, depending on the thresholds used to identify UCEs and tuning parameters of the HMM used to identify CNEEs, some UCEs will not be found in the set of CNEEs identified; this frequently happens when the core regions of UCEs are short, resulting in non-significant log-odds scores, which depend on length, when comparing the likelihood of CNEE sequences on trees with ‘neutral’ or ‘conserved’ branch lengths, the two states often used in the HMM. Finally, whereas UCEs are by definition found throughout the clade of interest (e.g., vertebrates) -- widespread presence across a clade is one criterion for inclusion in this marker class -- CNEEs can appear at variable nodes in the Tree of Life; the appearance and disappearance of CNEEs has been studied to gain insight into the origin of phenotypic traits (e.g., Lowe et al. 2011; Lowe et al. 2014).

In this study, we compare the phylogenetic performance and evolutionary dynamics of three classes of noncoding genomic markers: CNEEs, UCEs, and introns. We focus on noncoding regions because they appear to be promising for vertebrate phylogenetics, and we agree with suggestions that transcriptomes may have undesirable phylogenetic properties, especially at high taxonomic levels. Our questions in analyzing CNEEs in a phylogenetic context include: do CNEEs resolve phylogenetic relationships as well as UCEs or introns? How do the substitution dynamics of CNEEs compare with those of UCEs and introns? Do CNEEs exhibit alignment and evolutionary properties that are desirable for phylogenomic analysis? And finally, how easily can CNEEs be accessed in non-model species, and what sorts of protocols are recommended for their large-scale deployment in phylogenomics?

## Methods

### Compiling CNEEs in Genomic Data

We previously explored the use of CNEEs as markers of regulatory evolution in vertebrates (Lowe et al. 2014). As part of an effort to understand the genomic basis of feather evolution using CNEEs, we first aligned 19 vertebrate genomes using BlastZ, MultiZ and chaining of local alignments (Kent et al. 2003; Schwartz et al. 2003; Blanchette et al. 2004). We then used a hidden a phylogenetic hidden Markov model (HMM, Siepel and Haussler 2004) to determine regions of the genome (both coding and noncoding) that evolved slower than a benchmark set of 4-fold degenerate sites. The phylogeny of the 19 vertebrates used by Lowe et al. (2014) is well known and was assumed as fixed for all genes and genomic regions prior to analysis. This assumption is standard in pipelines for identifying CNEEs; although it ignores the possibility of incomplete lineage sorting (ILS), discordance due to ILS between the local genomic region and the vertebrate species tree we assumed, which has relatively long branches, is likely to be rare. Still, the potential biases incurred by assuming a fixed tree while identifying CNEEs should be explored, since ILS is known to influence parameter estimates of other macroevolutionary phenomena, such as molecular clocks, substitution rates and reconstruction of ancestral sequences (Burbrink and Pyron 2011; Groussin et al. 2015; Mendes and Hahn 2016). Branch lengths of the tree for the neutral class of sites were determined using maximum likelihood to find optimal branch lengths for the set of 4-fold degenerate sites. Conserved sites were defined as those exhibiting a better fit to a tree with branches no greater than 0.3 (30%) of the length of the 4-fold degnereate tree. The HMM had two states, “conserved” and “neutral” and the tuning parameters for the transition rate between states in the HMM were set with an expectation that CNEEs would on average have a length of 45 bp. This protocol yielded a total of 957,409 conserved elements in total, of which 605,756 fulfilled the criteria for a CNEE. Whereas Lowe et al. (2014) used 602,539 CNEEs in their study, we retained 3207 CNEEs that were discarded in Lowe et al. (2014) because they were not assigned to chromosomes in the chicken assembly used (galGal3), making for a starting total of 605,756 CNEEs. For a detailed account of the bioinformatics pipeline by which we initially determined a working set of CNEEs, see Lowe et al. (2014). Candidate CNEEs were filtered from this vertebrate-wide set of 605,746 elements referenced on chicken to retain loci ≥ 400 bp in length (n = 6182). We focused on CNEEs ≥ 400 bp long so as to use a set of loci expected to contain at least a moderate number of variable and parsimony-informative sites.

We then chose 14 exemplar species from the Avian Phylogenomics Project (Jarvis et al. 2014), including chicken, as a test case for phylogenomic analysis (see Supplementary File S1). These species were chosen so as to capture major branches of the avian tree as it is now known, and in some cases pairs of species were chosen to determine if our analyses could recapitulate known or expected relationships (e.g., flamingo and grebe, penguin and loon). This group of 14 species also contains clades that are still unresolved or contentious, such as the precise order of the multiple outgroups to passerine birds (Hackett et al. 2008; Jarvis et al. 2014; Prum et al. 2015). We also included data from draft genomes of an emu (*Dromaius novaehollandiae*) and Chilean Tinamou (*Nothoprocta perdicaria*) from Baker et al. (2014) so as to explore the hypothesis of ratite paraphyly (Harshman et al. 2008; Phillips et al. 2010; Smith et al. 2013), to make a total of 16 ingroup species. Using an American Alligator (Alligator mississippiensis, Green et al. 2014) genome as an outgroup sequence brought the total taxa used to 17 (SupplementaryFile S1). Blastn searches with chicken query CNEE sequences were used against each of the 16 non-chicken target genomes at an e-value cutoff 1e^-10^. CNEEs with no missing species were retained (n = 3822), and *de novo* aligned with default global alignment parameters in MAFFT v. 7.245 (Katoh and Standley 2013).

Intron alignments were assembled from the Avian Phylogenomics Project data of Jarvis et al. (2014); however, individual introns were used rather than alignments concatenating introns within each protein-coding gene. The SATé-MAFFT alignments provided by Jarvis et al. (2014) were reduced to the taxon subset of interest and gap-only columns removed. Loci greater than 400 bp in aligned sequence length, including the alligator outgroup sequence and with no more than 3 missing species were retained (n = 3733). It is noteworthy that it was straightforward to compile ~3700 fully populated CNEE alignments of 400 bp or greater, whereas there were only 998 (26.7%) fully-populated orthologous introns from birds available; we will return to this point in the discussion. Orthologous sequences from Emu and Chilean Tinamou were identified with blastn searches against draft genome assemblies for these species with chicken, ostrich, and white-throated tinamou queries, and were profile-aligned to the existing Jarvis et al. (2014) alignment with MAFFT. Because SATé-MAFFT yields relatively gappy alignments that are nonetheless “better” than MAFFT-only alignments by some optimality criteria (B. Faircloth, pers. comm.), comparing alignment statistics using SATé-MAFFT and MAFFT may bias the results. We therefore applied both SATé-MAFFT and MAFFT to all three marker types to enable side-by-side comparisons. For the CNEE alignments, we recapitulated the precise SATé-MAFFT alignment protocol of Jarvis et al. {, 2014 #4295), including post-alignment trimming with their custom python script ‘filter_alignment_fasta_v1.3B.pl’, except that we used SATe v. 2.2.7 with MAFFT v. 6.717 (Jarvis et al. used SATé 2.1.0 and MAFFT 6.860b). Ultraconserved elements (UCEs: n = 3679, representing the full set from Jarvis et al. 2014) were compiled as described for introns. There was a higher number of fully-populated alignments for UCEs of 400 bp or greater (n = 3669; 99.7%) than for introns.

As expected, there was overlap between our sets of CNEEs, UCEs and introns. For example, 1497 (39.2%) CNEEs overlapped at least one UCE. The degree of overlap between introns and the other two data sets was much lower: there were 6 introns overlapping CNEEs and 3 introns overlapping UCEs (both < 0.2%). Because UCE loci typically include the conserved core region in addition to the flanking regions, this overlap could lead to non-independence of our analyses. Therefore, in addition to analyzing the full set of UCEs and CNEEs, we also analyzed non-overlapping data sets of CNEEs and UCEs; in general, we found that our results held for overlapping and non-overlapping subsets of data, and we suggest that even if our CNEE and UCE data sets overlapped completely, analyzing just the core or flanking regions alone would help clarify the difference in dynamics and performance between these genomic regions.

### Measures of alignment quality and substitution dynamics

Alignment lengths, proportions of variable and parsimony informative sites, GC content, and the amount of missing data per alignment matrix (here defined as the number of gaps and uncalled bases per total cells in the nucleotide matrix) were calculated with AMAS (Borowiec 2016). Average pairwise nucleotide identity between species within each locus, and the proportion of gaps per base pair of aligned sequence were calculated with custom Perl scripts. Unlike the AMAS calculation of missing data, gaps per bp aligned considers only genuine gap characters (ignoring uncalled bases) and excludes leading and trailing gaps as well as gaps adjacent to uncalled bases; it is equivalent to internal gaps in the alignment per total called bases. TrimAl v. 1.2rev59 (Capella-Gutiérrez et al. 2009) was used for column-based alignment filtering, with the ‘automated1’ option to choose trimming parameters heuristically based on input alignment characteristics. We recognize that TrimAl and other alignment trimmers may not necessarily improve phylogenetic analysis in some cases (Tan et al. 2015), but we use them here strictly as a standard metric for comparing alignment “quality”, without subsequent phylogenetic analysis on the trimmed alignments. Additionally, we note that in many of the analyses in Tan et al. (2015), alignment trimmers performed marginally worse only under unsustainably high levels of trimming. Model-averaged transition/transversion rate ratios (Ti/Tv), the proportion of invariant sites when appropriate and the gamma shape parameter (α) were estimated for each alignment with jModelTest v. 2.1.7 (Darriba et al. 2012). jModelTest runs included six substitution models (JC, F81, K80, HKY, SYM, and GTR), with invariant sites and unequal base frequencies allowed and rate variation modeled with 4 gamma categories.

### Phylogenetic analyses and measures

RAxML v. 8.1.4 (Stamatakis 2014) was used to construct 200 bootstrap replicate gene trees from each unpartitioned alignment for each locus with a GTR + Γ substitution model; these were rooted with the Alligator outgroup in DendroPy v. 3.12.0 (Sukumaran and Holder 2010; Sukumaran and Holder 2015). MP-EST v. 1.5 (Liu et al. 2010) was used to infer species trees for each marker type from the input set of rooted RAxML bootstrap trees. Each analysis used three full MP-EST runs starting with a different random number seed and 10 independent tree searches within each run. Highest scoring trees from each search were used to build a majority-rule extended (MRE) consensus tree for each MP-EST run using RAxML. Per-site consistency indices (CI) were calculated with PAUP v. 4a149 (Swofford 2002) using the MRE consensus gene tree of the 200 RAxML bootstrap replicates for each locus. We did not compute consistency indices on species trees because gene tree heterogeneity can distort statistics like CI when all gene trees are forced onto a single topology (Mendes and Hahn 2016). Average bootstrap supports are also reported for MRE consensus gene trees.

### Phylogenomic subsampling

Phylogenomic subsampling (Edwards 2016) was used to assess the stability of specific clades for different subsets of each of the CNEE, intron and UCE data sets. Data sets of increasing numbers of loci (n = 50, 100, 200, 300, 400, 500, 1000, 1500, 2000, 2500, 3000, and 3500 loci) were built by sampling loci with replacement from within each marker type, and repeating the process to generate 10 independent replicates of a given number of loci within each marker type. MP-EST was then run on each of the 10 replicates as described above, except that only a single MP-EST run (but with 10 independent tree searches) was performed for each replicate. Summary measures are reported by counting the frequency of splits from among the set of MP-EST output trees for each replicate rather than from a consensus tree.

## Results

### Alignment and variability metrics for non-coding markers in birds

#### Alignment lengths and variability

Fig. 2 shows the distribution of alignment lengths among the three marker types and the percentage of variable sites within each alignment. With the constraint that each alignment must equal or exceed 400 bp, introns had longer alignments (up to 22,138 bp) than CNEEs (longest alignment, 1829 bp; Fig. 2a-c). UCE alignments based on those of Jarvis et al. (2014) varied from 2,126 – 4279 bp. CNEE alignments exhibit a higher fraction of populated bases per alignment than do introns and UCEs, with 1210 out of 3822 CNEE alignments (31.7%) possessing >99 % of populated bases (Fig. 3a and b). No intron alignments and only a single UCE alignment possessed this high a nucleotide matrix occupancy, whether considering any undetermined base or gaps between called sequence alone (Fig. 3c). CNEEs also exhibited a much lower percent of each alignment that was deemed low quality by trimAl than did introns or UCEs (Fig. 3d, Supplementary File S2. Whereas 1003 out of 3822 CNEE alignments (26.2%) retained >99 % of bases after trimming, only 1 of the UCE alignments and none of the intron alignments retained this much after trimming (Fig. 3d). As expected, both introns and UCEs were more variable than CNEEs (Fig. 2d-f;Supplementary Fig. S1a). The number of parsimony informative sites per alignment varied among markers in a similar way, with CNEEs having the fewest and introns having the most (Supplementary Fig. S1b). The number of variable sites scaled more linearly with alignment length for introns (r = 0.992, P < 0.00001) than for UCEs (r = 0.666, P < 0.0001) or CNEEs (r = 0.228, P < 0.00001; Fig. 2d-f). Although the alignment and variability statistics for UCEs changed significantly when analyzed using the MAFFT-only pipeline we used for CNEEs, the magnitude of the differences were small and trends among markers did not change (Supplementary File S3). Similarly, when we re-aligned all three marker types with the SATé-MAFFT used by Jarvis et al. (2014), overall trends and differences between markers were unchanged (Supplementary File S4).

**Figure 2:**
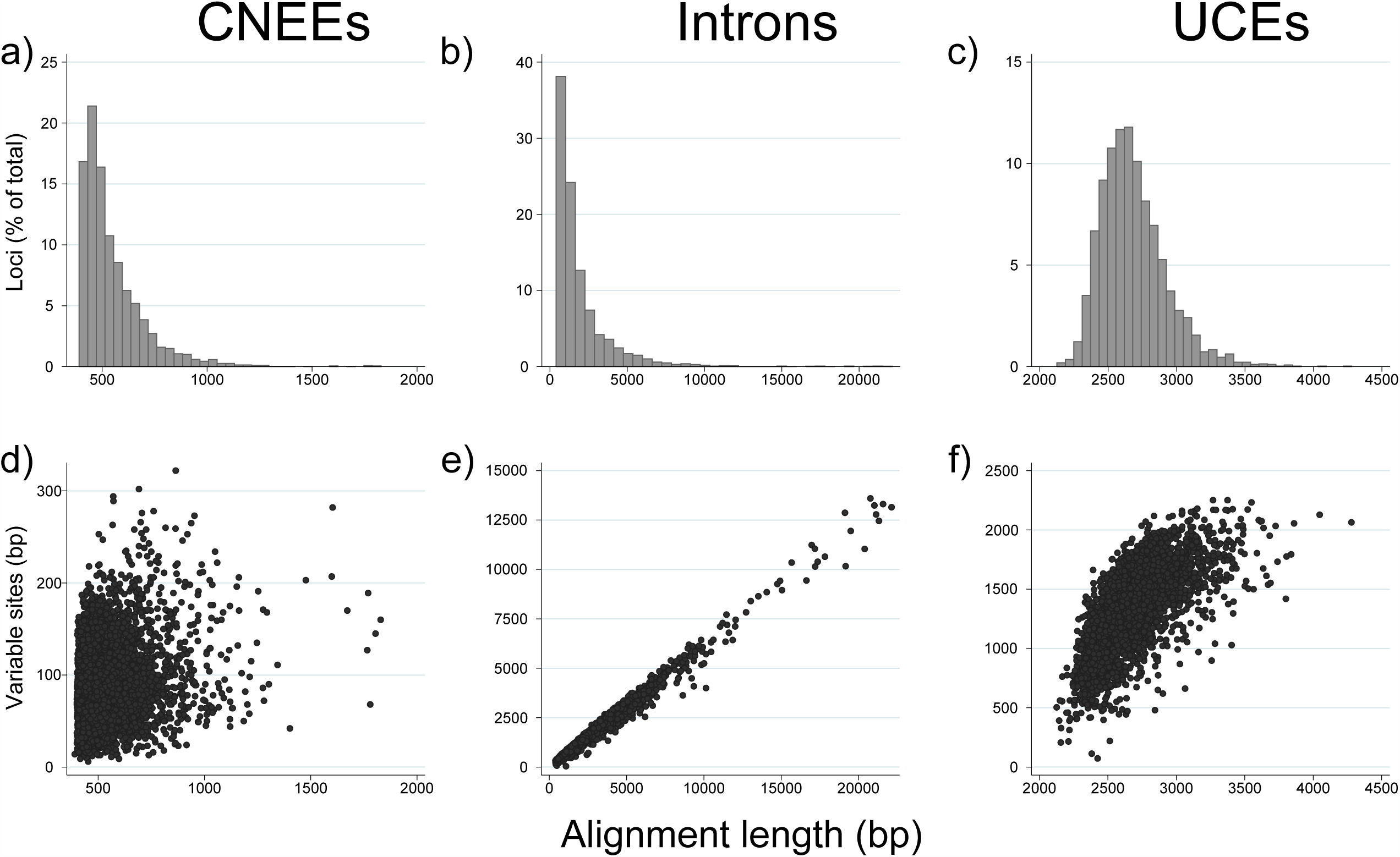
*Top row:* Distribution of aligned sequence lengths for a) CNEEs (3822 loci), b) introns (3579 loci), and c) UCEs (3679 loci). *Bottom* row: Correlations between alignment length and number of variable sites for d) CNEEs (r= 0.2277, P< 0.00001), e) introns (r= 0.9918, P< 0.00001) and f) UCEs (r= 0.6665, P< 0.0001).

**Figure 3.**
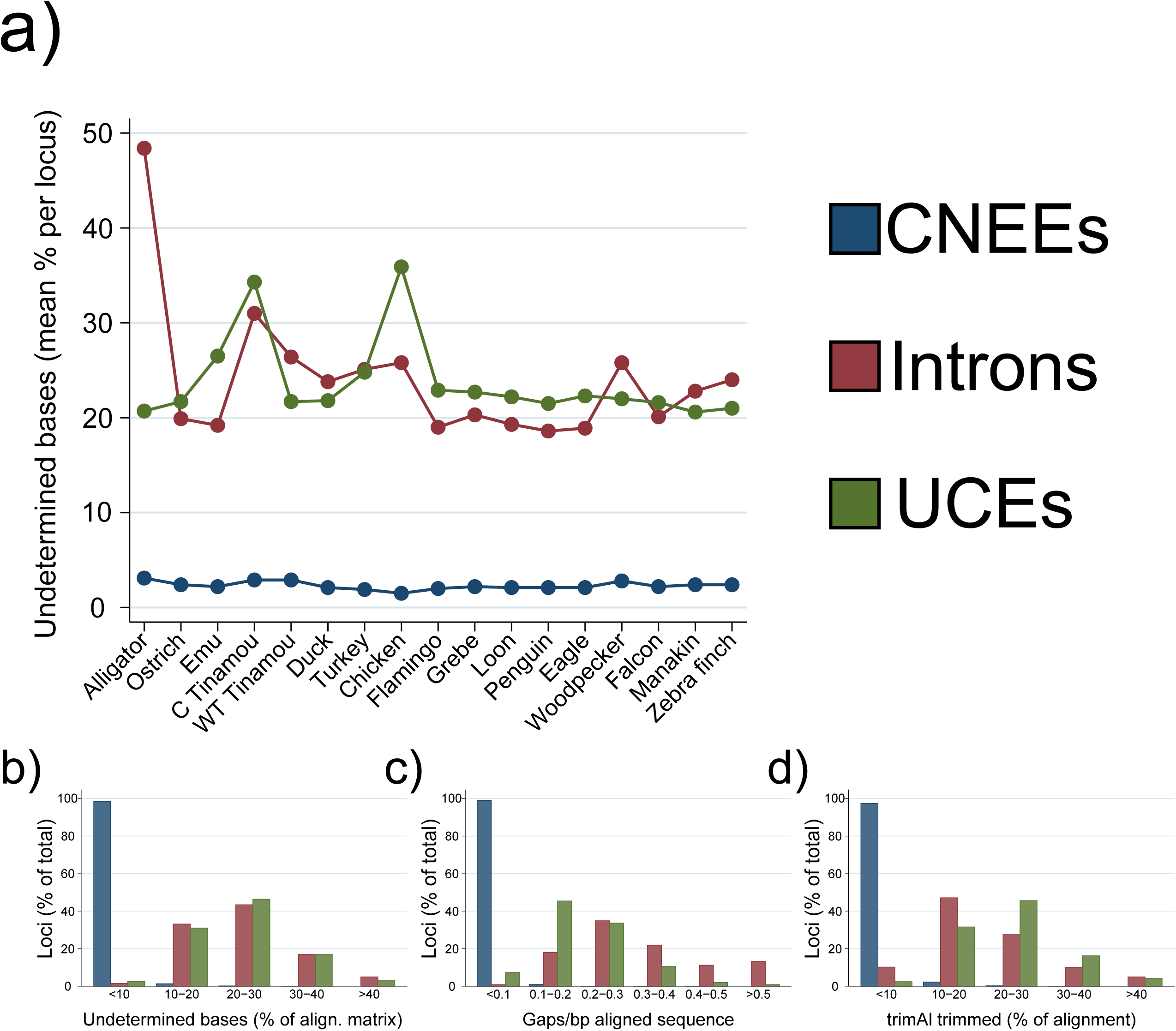
Variation in alignment gappiness among marker types. a) Average percentage of undetermined bases per alignment for each marker type and taxon. Here, undetermined bases indicates both gaps and Ns in each alignment. Alignments that were missing any species were excluded before analysis. b) Same data as in a, expressed as a histogram. c) Distribution of gaps per aligned base pairs, here including only genuine missing sequence. d) Distribution of the percentage of each alignment remaining after trimming with TrimAl. See methods for further discussion.

#### GC-content and substitution dynamics of noncoding markers

CNEEs exhibited systematically lower GC-contents than did introns or UCEs (Fig. 4a and b). There was a correlation between the GC-contents of different noncoding markers across species, presumably indicating a genomewide effect on base composition that influences all three marker types (Supplementary File S5). A notable outlier in GC-content across all three marker types is the Downy Woodpecker *(Picoides pubescens),* with average GC contents of 37.52% (CNEEs), 42.44% (introns) and 40.48% (UCEs), values that deviate from the grand mean for each marker type often 10 times more than for other species (Fig. 4a and b). High variance in GC-content can complicate phylogenomic analyses, since most phylogenetic models assume that all species in the analysis share a similar equilibrium base composition (Lockhart et al. 1994; Foster and Hickey 1999; Mooers and Holmes 2000). We found that the variance in GC-content among species was lowest for CNEE markers (average variance = 0.82), and higher for intron and UCE markers (average variance = 5.76 and 3.91, respectively; Fig. 4c). These substitution dynamics held in non-overlapping sets of CNEEs and UCEs (Supplementary File S6).

**Figure 4.**
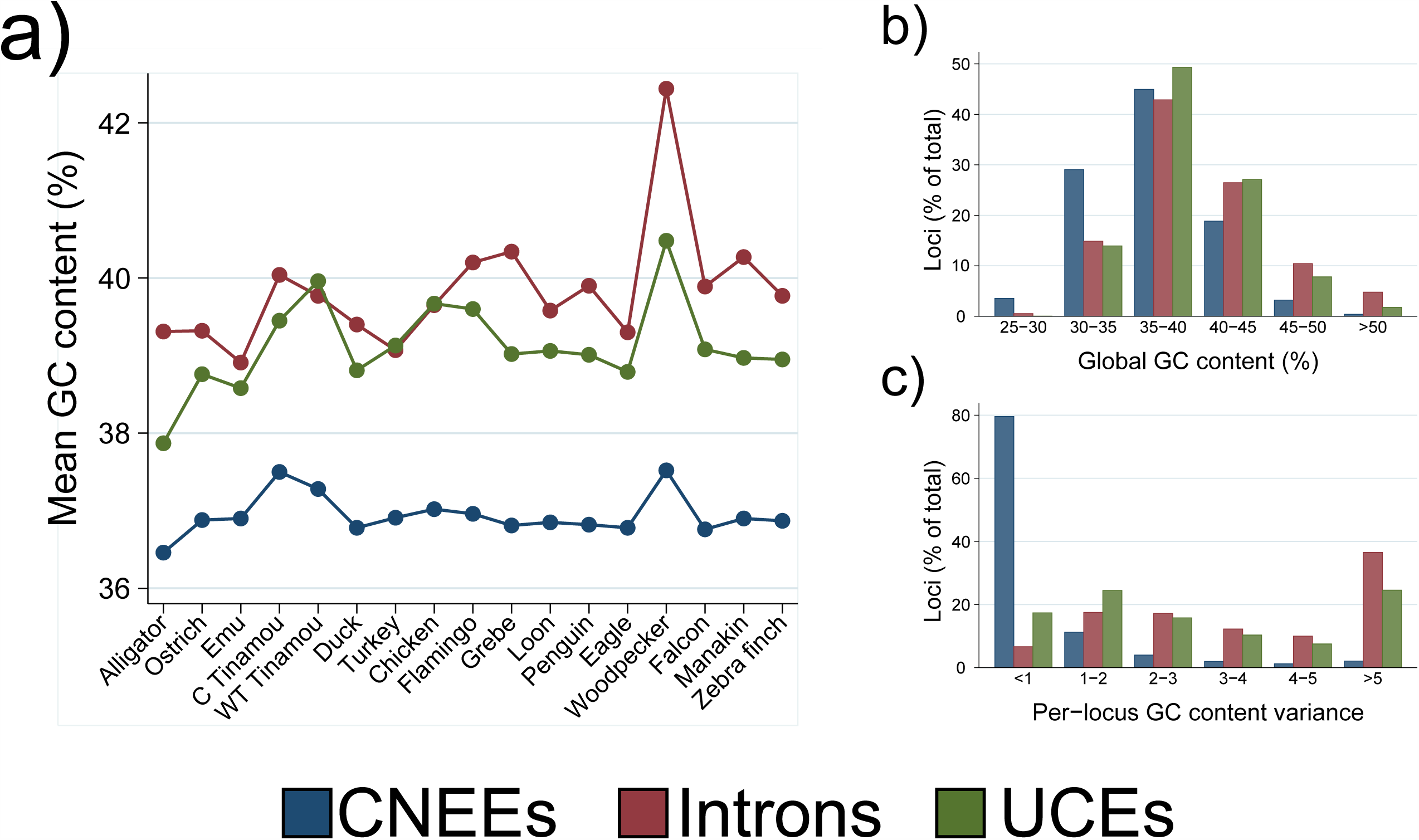
Patterns of GC content variation among markers and taxa. Alignments in which a taxon is entirely absent are omitted from all calculations. a) Per-taxon values for mean GC content for each marker type. b) Distribution of GC content among species for each marker type. c) Variance in GC content among taxa for each marker type.

Using jModelTest, we evaluated the substitution dynamics and optimal substitution model for each alignment. On average, CNEEs exhibited higher transition/transversion rate ratios (average 2.44) than did introns (1.90) or UCEs (1.79; Supplementary Fig. S1d). CNEEs also exhibited intermediate estimates of the gamma shape parameter (average 1.46) compared to introns (7.81) or UCEs (0.92; Supplementary Fig. S1). Overall, although all three markers displayed a similar range of nucleotide substitution models, the most complex models (GTR+ G+ I and GTR+ G) were least prevalent as the best-fitting model for CNEEs (7.2 and 29.2% of loci, respectively) than for introns (13. 7 and 73.2%) or UCEs (74.7 and 23.6%; Supplementary File S7). CNEEs displayed significantly higher consistency indices (mean = 0.92 for full and non-overlapping set) than UCEs (mean = 0.82; p < 0.00001) or introns (mean = 0.82; p < 0.00001); Supplementary Fig. S1, Supplementary File S6).

### Phylogenomic signal and consistency of noncoding sequences

As expected from the rank order of variability of each of the three marker types, gene trees made from CNEE alignments exhibited the lowest average bootstrap support, with introns and UCEs having progressively higher support (Supplementary Fig. S1c). However, the estimates of overall phylogenetic relationships and clade support as judged by species tree analyses were generally concordant among marker types and with previous analyses using larger data sets (Jarvis et al. 2014). All markers recovered ratite paraphyly, with the emu clustering with the two tinamous to the exclusion of the ostrich at 100% bootstrap support (Fig. 5a-c). In all three trees, the Neognathae are monophyletic and the three taxa representing Galloanseriformes (Chicken, Turkey and Peking Duck) were monophyletic at 100%, appearing as expected as sister to all the remaning taxa (Neoaves). All branches in the MP-EST species trees in this study achieved ≥ 95% for all marker types, except for two branches in the total CNEE tree, two branches in the total intron tree, and one branch in the total UCE tree. The branches in question invariably involved relationships among the outgroups to passerine birds and falcons, a clade termed Australaves (Jarvis et al. 2014; Prum et al. 2015, Fig. 5d-f). Whereas the total CNEE tree suggests that the Bald Eagle is closer to this clade than the Downy Woodpecker (albeit with only 72% and 56% bootstrap support, respectively, for these two branchings), both the total intron and UCE trees support the reverse branching order, with first Downy Woodpecker (at 87% and 70% bootstrap support for introns and UCEs, respectively), then Bald Eagle (with 100% support in both cases) forming successive sister groups to the Australaves. Depending on how one likes to draw bootstrap support cutoffs in phylogenomics analyses, there is no case among the total marker trees of strongly supported conflict in overall species tree estimates among the three marker types for any cutoff greater than 87%. This trend largely held for phylogenetic analysis of the non-overlapping subsets of CNEEs and UCEs (Supplementary Fig. S2): support values increase for CNEEs (72% to 89% for eagle+falcon/passerines and 56% to 85% for woodpecker+other ‘land birds’), and decrease for UCEs (70% to 62% for woodpecker+falcon/passerines). When we confine phylogenetic analysis to the 1000 loci with the highest variability or most highly supported gene trees, the results are largely similar (Supplementary Fig. S3).

The relationships obtained for the three marker types are also similar to most of the analyses produced by the Avian Phylogenomics Project (Jarvis et al. 2014, Fig. 5d-f) and a recent, more taxon-rich tree for birds produced with 259 loci, most of which were derived from coding sequences (Prum et al. 2015). A source of disagreement for the taxa that we have sampled involved the sister group to Australaves (Prum et al. 2015). Although both Prum et al. (2015) and Jarvis et al (2014) generally produced trees placing Woodpeckers closer to Australaves than eagles, neither paper produced this result unambiguously; whereas Jarvis et al. (2014) achieved 100% support for a sister clade to Australaves that included both woodpeckers and eagles in their total evidence concatenation tree (TENT) tree using ExaML, other analyses from Jarvis et al. (2014), as well as the results of Prum et al., (2015), placed woodpeckers as sister to Australaves, with eagles falling outside this clade, albeit with highly varying levels of support. The relationships among waterbirds (penguin, loon, flamingo and grebe), although consistent across analyses and markers in this study, constitute another region of disagreement with studies employing more taxa. Whereas this study and Prum et al. (2015) suggest monophyly of the four water birds sampled here (Aequorlitornithes), many of the Jarvis analyses, including their TENT analysis, suggested paraphyly of this clade. Because our taxon sampling is so low we naturally defer to these larger studies for the provisionally ‘correct’ results for these clades, although we note that these larger studies were not able to robustly resolve all relationships, including the two clades discussed here.

**Figure 5.**
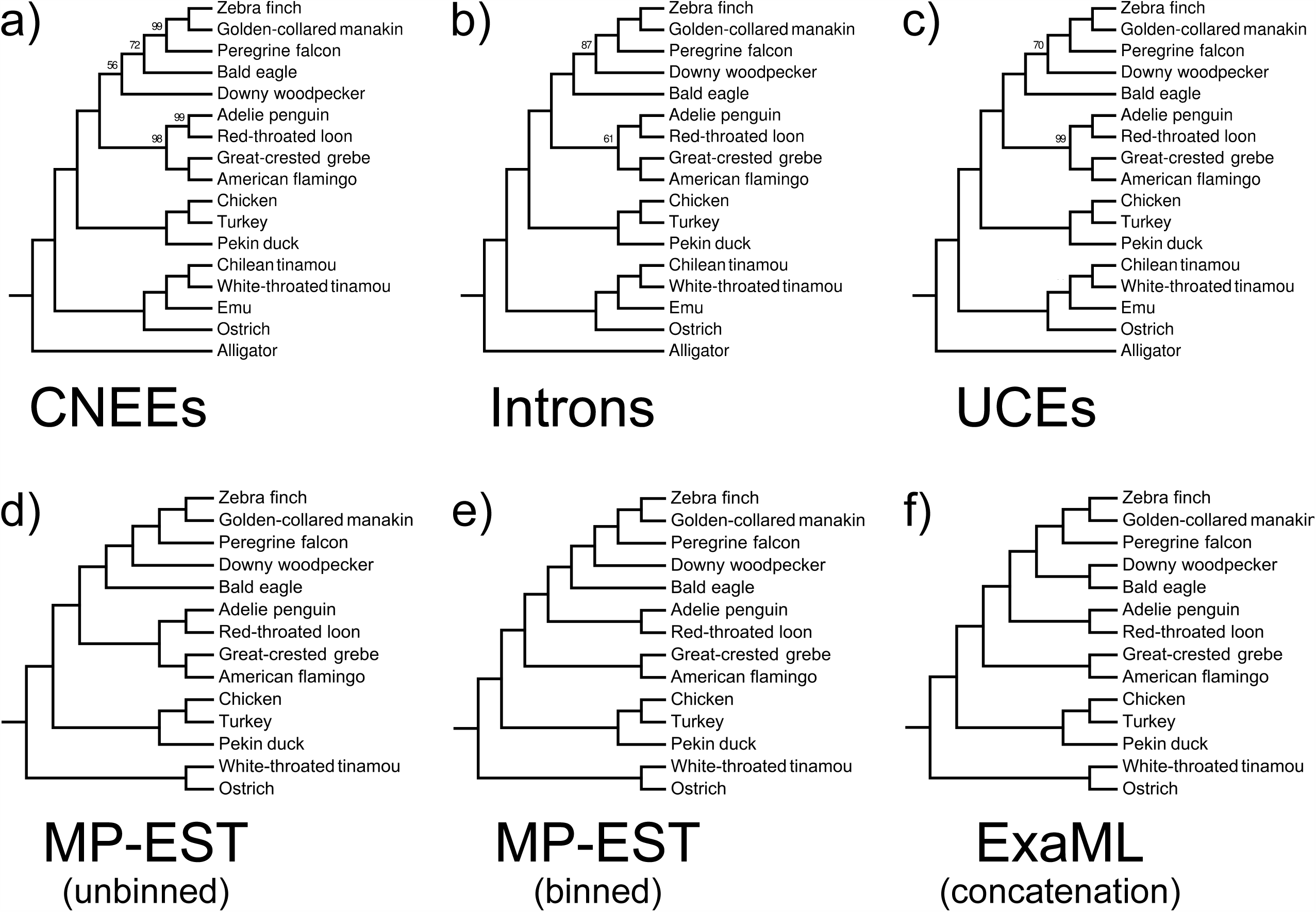
Species tree topologies discussed in this study. *Top row:* MP-EST species trees for a) CNEEs (3822 loci), b) introns (3579 loci), and c) UCEs (3679 loci), with support values < 100% indicated. *Bottom row:* Total evidence nucleotide trees (TENT), each built from 2516 introns, 3769 UCEs, and 8251 protein-coding genes, from Jarvis et al. (2014), pruned to the taxon set used in the current study. d) MP-EST unbinned analysis, e) MP-EST binned analysis and f) concatenated analysis. Support values are omitted from the pruned Jarvis et al. trees. The main tree presented in Prum et al. (2015) is identical to the tree depicted in panel d, assuming that Chilean Tinamou and Emu would fall where found in other studies.

We conducted phylogenomic subsampling to study the accumulation of signal as the number of loci increases for two expected clades that ultimately achieve high certainty for all data sets as well as for the two uncertain clades described above. The two high-confidence relationships we examined were the paraphyly of ratites and the sister group to passerines (i.e. falcons; Fig. 6a and b). We found that all three marker types established high confidence in the paraphyly of ratites by 200 genes, with introns accumulating signal somewhat faster than CNEEs and UCEs (Fig. 6a). By contrast, the falcon+passerine clade achieved consistent 100% support at 1000 loci for introns and UCEs, whereas CNEEs did not achieve an average of 100% support for the number of loci analyzed here, peaking at 98% support at 3500 loci and 99% with the full data set (Fig. 6b). For the monophyly of the waterbird clade (Fig. 6c), we found that the accumulation of signal was more rapid for CNEEs and UCEs, and less rapid for introns. Introns achieved an average bootstrap support of only ~70% for subsamples of 3500 loci (only 61% for the full data set), whereas average support of similarly sized subsamples of CNEEs and UCEs approached 100% (98 and 99%, respectively, for the full data set). For this clade, no marker type exhibited monotonically increasing average support with larger subsamples of loci, although the lack of monotonic increase was much more pronounced for introns than for the other two markers (Fig. 6c). The subsampling results for the sister to Australaves are more interesting, in so far as they begin to suggest genuine conflicts between the marker types. Whereas both introns and UCEs accumulate stronger signal favoring a woodpecker+Australaves clade (87 and 70%, respectively; Fig. 5b and c; Fig. 6d), the CNEEs instead accumulate stronger signal favoring an eagle+Australaves clade, approaching 72% (Fig. 5a, 6d). Whereas CNEEs exhibit a threshold of sorts for the accumulation of signal for the waterbird clade, increasing in average support and number of replicates achieving > 70% support at 500 loci (in part an artifact of the particular intervals chosen for subsampling), introns suggest a threshold at 1500 loci for the woodpecker/Australaves clade (Supplementary Fig. S4).

**Figure 6.**
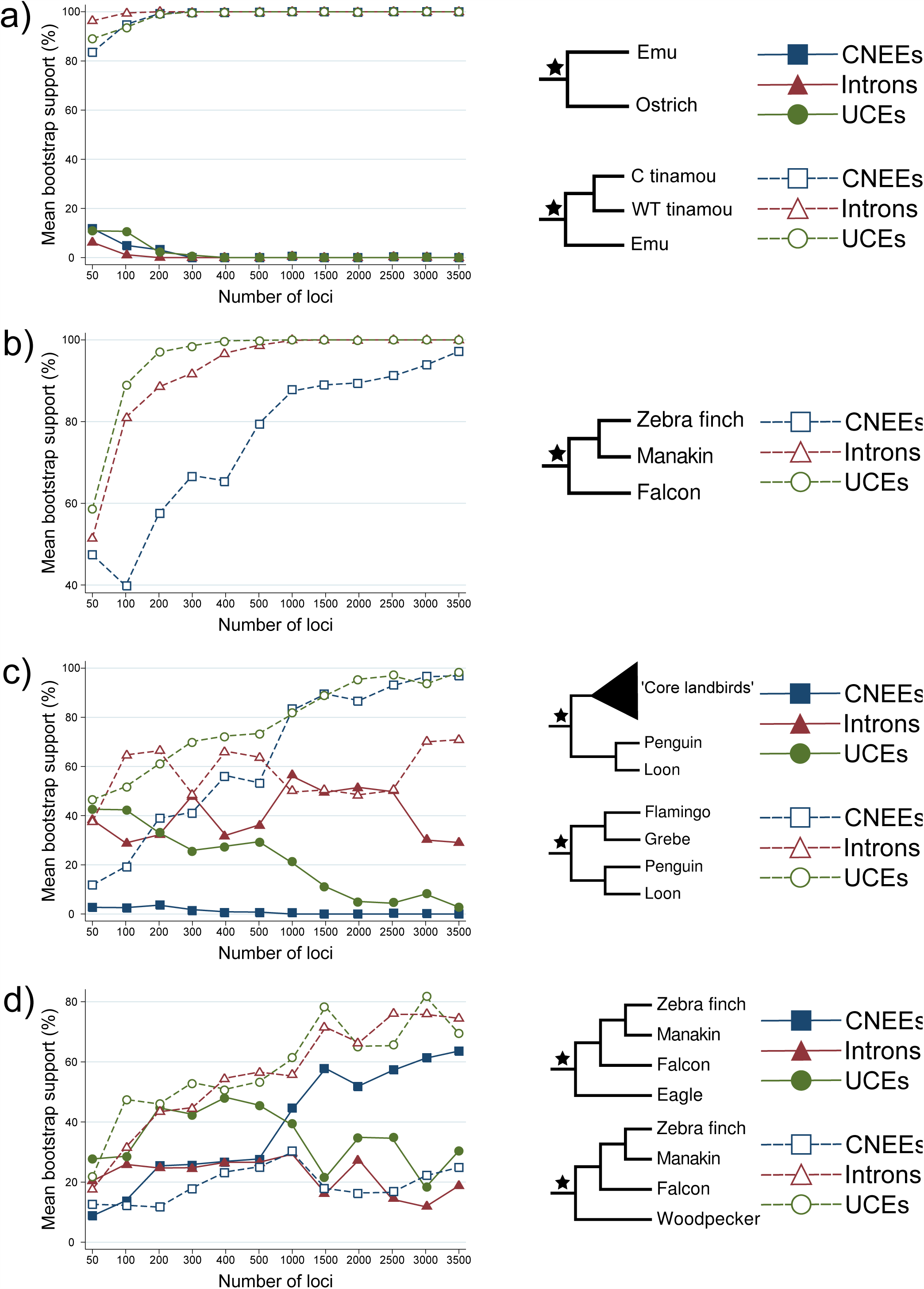
Phylogenomic subsampling, with MP-EST species trees inferred for twelve datasets of increasing numbers of randomly chosen loci, and with ten replicates per dataset. In each row, left panels plot the mean bootstrap support among the ten MP-EST replicates for each marker type and data set size. At right are the two branches, indicated by stars, whose support is investigated by subsampling, with open and solid markers indicating support for one or the other branch. a) Trends in support with increasing numbers of loci for paraphyly of ratites. b) Trends in support for the branch uniting falcons as the sister group to passerine birds. c) Trends in support for competing hypotheses placing either core landbirds (songbirds+falcon+eagle+woodpecker) or (flamingo+grebe) as the sister group to (penguin+loon). d) Trends in support for either bald eagle or downy woodpecker as the sister to (songbirds+falcon).

## Discussion

In this study we explored the evolution of CNEEs, a class of noncoding marker that has not received attention in terms of its utility for phylogenomics, and compared them to the performance of two other classes of noncoding markers, introns and UCEs. Overall, the full data set of CNEEs performed well compared to introns and UCEs, with a similar number (1-2) of branches in our 17-taxon tree achieving less than 95% support. The utility of CNEEs for phylogenomic analysis will depend somewhat on the values held by different researchers. If a researcher values high support for branches achieved quickly as numbers of loci are increased, at the expense of more uncertain and gappy alignments with missing species, then introns and UCEs clearly outperform CNEEs. However, if a researcher favors higher certainty and quality of alignment, and a better fit of alignments to the equilibrium assumptions of most phylogenetic models of nucleotide substitution, then CNEEs may offer advantages. The major advantages of CNEEs are the ease of obtaining large numbers of high-quality alignments without missing species, their low homoplasy and their low variance in GC across species. Despite their low variability, and the correspondingly weak support in gene trees, CNEEs produced a species tree that rivaled those produced by a similar number of intron and UCE alignments (Fig. 6). Indeed, given that the CNEE alignments were the shortest of the three marker types, one could argue for the overall efficiency of CNEEs in terms of phylogenetic resolution per base pair sequenced as compared to the other two markers.

A critical factor is the number of markers of reasonable length available to phylogeneticists, and the ease of producing fully-populated alignments, since these factors could place a limit on phylogenomic resolution of a particular marker type. Introns are numerous in vertebrate genomes, on the order of several times the number of genes, which usually number about 15,000-20,000. However, orthologous introns often vary substantially in length among taxa (Vinogradov 2002; Waltari and Edwards 2002; Pozzoli et al. 2007; Zhang and Edwards 2012); due to their high variability and length differences, gaps will be frequent, with many alignments > 400 bp having large numbers of unfilled (missing) bases. Conserved elements are very numerous in vertebrate genomes, with as many as 3.6 million elements detected in mammals, over 80% of which are noncoding (Lindblad-Toh et al. 2011). However, the average length of these elements is often < 50 bp. Faircloth et al. (2012) were able to assemble 5599 unique UCEs, which need to achieve a certain minimum length of the core region to be detectable by hybrid capture methods. Bejerano et al. (2004) found only 481 fully conserved UCEs longer than 200 bp in vertebrate genomes, and the total number of UCEs > 100 bp in vertebrates is estimated to be ~14,000 (Stephen et al. 2008). Today, markers designated as “UCEs” often contain loci that are not strictly UCEs, but rather CNEEs, whose core is often more variable than the original definition of UCEs (Bejerano et al. 2004). In this respect, the number of “UCE” loci has increased in recent studies and can overlap even more with loci designated here as CNEE.

It was straightforward to compile a data set of several thousand CNEE markers > 400 bp which contained all species, and most of which contained < 2 alignment indels. By contrast, because of their intrinsic variability or reliance on variable flanking regions, both introns and UCEs had between 20-40% missing data (gaps) per species per alignment, a consequence of their high indel rate, and it was challenging to find intron alignments that contained all 17 species in our study. These trends were evident despite the fact that all three marker types were harvested from whole genomes, as opposed to being generated using molecular methods such as hybrid capture. The reasons for the lower incidence of fully-populated alignments for introns does not seem to lie in the lower coverage of some of the genomes used since the CNEEs were harvested from the same source data. Rather, it seems to lie in the greater challenges of detecting introns via blast, or by challenges with genome annotations, or the great length of many introns, which undercuts search algorithms.

The three marker types exhibited differing patterns of nucleotide substitution, which could influence their phylogenetic performance, and which appear to be driven in part by overall levels of variation. For example CNEEs had the lowest level of among-lineage variation in GC content, a trait that conforms well with the equilibrium assumptions of most models of nucleotide substitution. To our knowledge we are the first to report the anomalous GC content of the Downy Woodpeker genome. It is likely that the higher fraction of transposable elements in this genome (~22%) compared to other birds investigated thus far, as reported by (Zhang et al. 2014), is linked to the outlier status of the Downy Woodpecker in terms of GC content, although we have not verified the prediction that TEs in this genome are higher in GC than other genomic regions. As expected due to their inclusion of the slowly evolving core as well as more rapidly evolving flanking sequences, UCEs exhibited high levels of among-site rate variation (low α) compared to introns and CNEEs. Although not necessarily detrimental to phylogenetic analysis, it is widely acknowledged that high levels of among site rate variation are more difficult to model than low levels (Vogler et al. 2005; Marshall et al. 2006; Holland et al. 2013). On the other hand, CNEEs exhibited the highest transition/transversion ratio among the markers; although high ts/tv ratios, like among-site rate variation, often lead to homoplasy, the higher consistency index among CNEEs appears here to be driven more by their low substitution rate than ts/tv ratio. CNEEs were markedly more AT-rich than the other two classes of markers, which, as a group, tend to be more AT-rich than coding regions. Although AT- versus GC- rich markers do not present any obvious advantages, Romiguier et al. (2013) recently suggested that, in mammals, GC-rich markers result in higher gene tree heterogeneity than AT-rich markers, possibly due to biased gene conversion, making phylogenetic analysis more challenging.

### Information Content of CNEEs for Phylogenetic Analysis

Overall, we found that CNEEs delivered an estimate of phylogenetic relationships that was as strong as that for UCEs and introns. For some expected phylogenetic results, such as the paraphyly of the ratites, the approach to phylogentic “certainty” (100% bootstrap support) was as fast as that for the other two markers. However, for other questions that appear to be gaining consistent support among phylogenomic data sets, such as the falconid sister group of the passerine birds, the approach to phylogenomic resolution was markedly slower than for UCEs or introns. And yet for other clades, such as the sister relationship between penguin/loon and flamingo/grebe, it was introns that failed to achieve high resolution compared to CNEEs and UCEs. Finally, CNEEs suggested a different sister group to falconids and passerines, namely eagles, at fairly high (~80%) and increasing support as more loci were accumulated, as compared to introns and UCEs, which favored woodpeckers as the sister group, again with high support. This result was the only case of moderately strong conflict among markers in our data set, and in our view, either result is plausible, given that this node was not resolved with certainty among larger data sets (Jarvis et al. 2014). For example, the fact that among the taxa we studied the woodpecker is a base compositional outlier more strongly for introns and UCEs than for CNEEs could be driving this difference in result. We were able to achieve high and consistent confidence for nearly all branches in our analysis without binning (Mirarab et al. 2014), suggesting that large numbers of loci, rather than concatenation of loci, remains a plausible way forward for phylogenomics (Liu and Edwards 2015). Although our results point to possible differences in performance and useful trends among these markers, because we only sampled 16 ingroup bird species as exemplars, the generality of these trends requires further investigation.

Finally, we do not expect CNEEs to provide resolution at low taxonomic levels or in phylogeography, where UCEs appear useful (Lemmon and Lemmon 2012; McCormack et al. 2013; Smith et al. 2014; Hamilton et al. 2016; Manthey et al. 2016), in part due to their low variability but also because, like UCEs, they are likely to be under strong background selection (Katzman et al. 2007).

In summary, CNEEs appear to be a promising tool for phylogenomic research. Their low variability compared to introns and UCEs is offset by the larger numbers of moderately long and high quality alignments that can be gathered from whole-genome data sets. In the future, as whole genomes become more readily available, phylogenomic data sets will increasingly be generated via statistical tools or extraction of large sets of alignments from aligned or unaligned genomes (Costa et al. 2016), rather than directly by wet lab bench work. Until that time arrives, wet-lab approaches to gathering loci, such as hybrid capture, will continue to be used. In either scenario, CNEEs should fare well, because they are readily identified by statistical means from whole genomes, and yet they would also be amenable to hybrid capture approaches. We expect that mixtures of noncoding phylogenomic markers, including CNEEs, will be helpful in understanding the dynamics of currently popular markers such as UCEs and introns and will contribute to resolving the Tree of Life.

## Supplementary Information

**Supplementary File S1.** Sources of noncoding markers used in this paper, including 14 exemplar species from the Avian Phylogenomics Project.

**Supplementary File S2.** Alignment summary measures for all loci.

**Supplementary File S3.** Comparison of UCE alignment and variability metrics for SATé- MAFFT alignments with emu and Chilean tinamou profile aligned, and for de novo alignment of all sequences with MAFFT.

**Supplementary File S4.** Comparison of alignment and variability metrics for de novo alignment with MAFFT (CNEEs) and SATé-MAFFT alignment with emu and Chilean tinamou profile aligned (introns and UCEs) as used throughout the manuscript, versus alignment following the pipeline described in Jarvis et al. (2015).

**Supplementary File S5.** Pairwise correlations in per-taxon GC content between marker types.

**Supplementary File S6.** Substitution dynamics of non-overlapping sets of CNEEs and UCEs.

**Supplementary File S7.** Summary of jModelTest output for all loci.

**Supplementary Fig. S1**. Distribution of variable and parsimony informative sites, gene tree resolution, and substitution dynamics across marker sets.

**Supplementary Fig. S2.** Phylogenetic analysis of non-overlapping sets of CNEEs (n = 2318 loci), introns (n = 3685 loci) and UCEs (n = 2232 loci).

**Supplementary Fig. S3.** Phylogenetic analysis of the 1000 most strongly supported gene trees for each marker (top row) and the 1000 most variable markers (bottom row).

**Supplementary Fig. S4.** Threshold analysis for subsampling across markers.

## Acknowledgements

We thank Brant Faircloth for help in assembling the UCE data set, and Brant Faircloth and Craig Lowe for helpful discussion and comments on the manuscript. This research was generously supported by the Harvard University FAS Research Computing and the Odyssey Cluster.

